# Establishing elements of a synthetic biology platform for Vaccinia virus production: BioBrick™ design, serum-free virus production and microcarrier-based cultivation of CV-1 cells

**DOI:** 10.1101/042275

**Authors:** Shuchang Liu, Ludmila Ruban, Yaohe Wang, Yuhong Zhou, Darren N. Nesbeth

**Affiliations:** Department of Biochemical Engineering, University College London, Bernard Katz Building, London WC1E 6BT, UK; Centre for Molecular Oncology & Imaging, Institute of Cancer, St Bartholomew’s and the London School of Medicine and Dentistry, Queen Mary University of London, London EC1M 6BQ, UK

**Keywords:** vaccinia virus, microcarrier, titre, virus, synthetic biology, serum free medium, bioprocess development, BioBrick™

## Abstract

Vaccinia virus (VACV) is an established tool for vaccination and is beginning to prove effective as an oncolytic agent. Industrial production of VACV stands to benefit in future from advances made by synthetic biology in genome engineering and standardisation. As a step toward realising these benefits we analysed the Lister Vaccinia virus genome with respect to refactoring options and propose a VACV genome engineering BioBrick™. We then used the CV-1 cell line to produce a conventional recombinant Lister strain VACV, VACVL-15 RFP in a serum-free process. CV-1 cells grown in 5% foetal bovine serum (FBS) Dulbecco’s Modified Eagle Medium (DMEM) were adapted to growth in OptiPRO and VP-SFM brands of serum-free media. Specific growth rates of 0.047 h^−1^ and 0.044 h^−1^ were observed for cells adapted to OptiPRO and VP-SFM respectively, compared to 0.035 h^−1^ in 5% FBS DMEM. Cells adapted to OptiPRO and to 5% FBS DMEM achieved recovery ratios of over 96%, an indication of their robustness to cryopreservation. Cells adapted to VP-SFM showed a recovery ratio of 82%. VACV production in static culture, measured as plaque forming units (PFU) per propagator cell, was 75 PFU/cell for cells in 5% FBS DMEM. VP-SFM and OptiPRO adaptation increased VACV production to 150 PFU/cell and 350 PFU/cell respectively. Boosted PFU/cell from OptiPRO-adapted cells persisted when 5% FBS DMEM or OptiPRO medium was present during the infection step and when titre was measured using cells adapted to 5% FBS DMEM or OptiPRO medium. Finally, OptiPRO-adapted CV-1 cells were successfully cultivated using Cytodex-1 microcarriers to inform future scale up studies.

## Introduction

Vaccinia virus (VACV) is an enveloped, brick-shaped particle typically 300×240×120 nm containing a double stranded DNA genome which, for the Lister strain (Garcel et al. 2007), is 189.4 kilobase-pairs (kb) in size, encoding up to 201 open reading frames (ORFs). VACV is arguably one of the most effective biotechnological tools in history by dint of the fact human antibodies raised against VACV tend also to recognise smallpox epitopes. VACV was used to eradicate smallpox via a global vaccination programme carried out by the World Health Organization (WHO) between 1966 and 1980 (Fenner et al., 1988).

VACV has also been used as a molecular biology tool to effect high-level transgene expression in mammalian cells, due in part to its ≈25kb capacity for accommodating recombinant DNA (Mackett and Smith, 1986; Hruby, 1990; Guse et al., 2011). Molecular biology techniques developed in this area have also enabled construction of a wide range of recombinant VACV vaccines in which selected epitopes or payloads are defined by recombinant DNA.

Recombinant VACV has been developed as an effective live vaccine against viral, bacterial and oncological diseases (Hruby, 1990; Lee et al., 1994; Timiryasova et al., 1999; Zhang et al., 2007), due to its ability to elicit vigorous antibody and T-cell mediated responses. Hiley et al. (2010) and Tysome et al. (2011) have also demonstrated the effectiveness of recombinant Lister strain VACV in targeting hypoxic tumours in human head and neck cancer.

Mass production of VACV for smallpox eradication was achieved by harvesting virus from lesions brought about by infection of live animals (Fenner et al., 1988). However, this method brought significant risk of contamination with microbiological agents and was superseded by viral propagation in embryonated hen eggs (EMEA 2012). Monath et al. (2004) investigated the use of MRC-5 cells to produce the New York City Board of Health (NYCBH) VACV strain for a Phase I clinical trial as a smallpox vaccine. As a human diploid cell line, MRC-5 has a finite in vitro life span that limits capacity for long-term cultivation and large-scale VACV production using diploid cell lines can be difficult as such cells typically do not grow well on microcarriers (Barrett et al., 2009).

Vero cells cultivated in serum-free media (Mayrhoffer et al. 2009) have been used previously to produce VACV and Monath et al. (2004) used Vero cells grown on microcarriers for VACV production at 1200L scale in bioreactors. In effect Vero cells are the workhorse of VACV production so in this study we sought to diversify the range of mammalian cell lines used for VACV production. CV-1 cells are an obvious first choice as they have already been used widely for VACV titration (Schweneker et al. 2012).

At laboratory-scale, scale-out strategies, such as roller bottles, T-flasks and the Nunc™ Cell Factory™, are commonly used to cultivate adherent cells for propagation of VACV. However, methods that can be scaled up, as opposed to scaled out, are the ideal solution for increasing the level of production, predictability and affordability for widespread application of VACV-based therapies. Toward this aim Bleckwenn et al. (2005) used HeLa S3 cells grown on microcarriers, at 1.5L scale, in a hollow fibre perfusion bioreactor setup to propagate VACV. Viral vaccine production in media supplemented with bovine serum has been discouraged by regulatory authorities such as the Food and Drug Administration (FDA), brings high variability between serum batches and can lead to variations in product yield and quality. Undefined components in serum may also provide a route for adventitious agent contamination. Bioprocesses that are serum-free and animal derived component free (ADCF) are now sought in order to reduce the contamination risk, ease the downstream processing artefacts and promote robustness and reliability for the production of VACV. Previous attempts to grow CV-1 cells in serum-free media (Steimer et al. 1981) replaced serum with other animal-derived products so did not remove routes for adventitious agent contamination.

Synthetic biology aims to render biological phenomena easier to engineer (Ye and Fussenegger 2014). An inevitable consequence of this aim is that biology becomes easier to manufacture. When applied to VACV production, and its exploitation in areas such as gene therapy and oncotherapeutics, synthetic biology offers the prospect of rapid design and assembly of viral payloads using interoperable tools, such as BioBrick™-formatted plasmids (Shetty et al., 2008), compatible with repositories containing thousands of components.

Synthetic DNA is now being used to construct large segments of eukaryotic genomes (Dymond et al. 2011) and construction of human artificial chromosomes (Kononenko et al. 2015) is now an established approach in gene therapy research. We feel it is then reasonable to expect that cells, or ‘chassis’, could in the near future be controlled by bespoke genomes designed for maximal VACV production yields and integration with scalable bioprocesses. Here we take steps toward such a synthetic biology platform for VACV production, firstly by proposing a BioBrick™-formatted plasmid for VACV genome engineering and appraising considerations for refactoring the VACV genome. Secondly, using a conventionally engineered VACV strain but a novel choice of propagator cell line, we retrofit VACV production from serum-containing media to serum-free media and test the resultant cells in terms of growth performance, viral productivity in T flasks and growth performance in a microcarrier-based cultivation platform.

## Material and Methods

### Cell Cultivation

CV-1 cells, product CCL-70™ from American Type Culture Collection (ATCC), were grown in High Glucose Dulbecco’s Modification of Eagle’s Medium (DMEM) from PAA Laboratories (Pasching, Austria), supplemented with 5% v/v foetal bovine serum (FBS) from batches A10409-1728 and A15112-2026 for three passages prior to this study. Cells were passaged twice weekly in T flasks and seeded at 1×10^4^ cells / cm^2^ for growth in 5% FBS DMEM and serum-free OptiPRO and 2×10^4^ cells/cm^2^ for growth in serum-free VP-SFM medium. Serum-free media was supplemented with GlutaMAX to 4mM and detached by treatment with TrypLE Select. All materials were sourced from Life Technologies, New York, USA, unless otherwise stated.

### Cell Banking and Revival

Cells adapted to growth in 5% FBS DMEM were cryopreserved in 90% FBS plus 10% v/v dimethyl sulfoxide (DMSO) from Sigma-Aldrich (Ayrshire, UK). Cells adapted to growth in serum-free media were frozen in a v/v mixture of; 45% fresh growth medium, 15% 2 day, 15% 3 day and 15% 4 day conditioned medium plus 10% DMSO and 0.1% v/v methylcellulose (Sigma). Cells were suspended in the cryopreservation medium at 3-5×10^6^ cells/mL and transferred to 2mL screw cap cryopreservation tubes (Eppendorf Ltd, Stevenage, UK) for storage in the liquid phase of a liquid nitrogen Dewar (Part No. 9902130, Statebourne Cryogenics, Tyne & Wear, UK). For revival, cryopreservation tubes were removed from liquid nitrogen and thawed in a SUB14 water bath at 37°C (Grant Instruments, Cambridge, UK). Upon thawing, cells in cryopreservation solution was diluted to 8mL and centrifuged at 1300rpm for 3minutes. The supernatant was withdrawn and cell pellet was resuspended in 8mL pre-warmed OptiPRO and transferred to a T 25 flask and incubated at 37°C, 5% CO_2_ in an MCO-19AIC incubator (Sanyo, Gunma, Japan).

### Counting cells cultivated using T flasks

Cells were detached from flask surfaces using standard trypsin treatment. Total cells in suspension were then counted using a TC10™ Automated Cell Counter (Bio-Rad, Hercules, USA) according to manufacturer’s instructions (document PN10016620 Rev B). Total viable cell counts in suspension were performed using standard trypan blue dye exclusion. Cells were stained with 0.4% trypan blue (#T8154, Sigma-Aldrich, Aryshire, UK) and counted using an Improved Neubauer haemocytometer (Hawksley, Lancing, UK) within three minutes of staining.

### Adaptation to Serum Free Media

Cells were grown in 10% FBS DMEM to a density of 1.3×10^5^ cells/cm^2^ in a T-25 flask (3.25×10^6^ cells total). Cells were then harvested into a total volume of 8mL growth media mix, containing, for each round of adaptation; 6mL, 4mL, 2mL, 0.8mL and finally zero mL 10% FBS DMEM made up to 8mL with serum-free media before further passaging. OptiPRO or VP-SFM brands of serum-free medium were used, as indicated in Figure 2.

**Figure 2.**
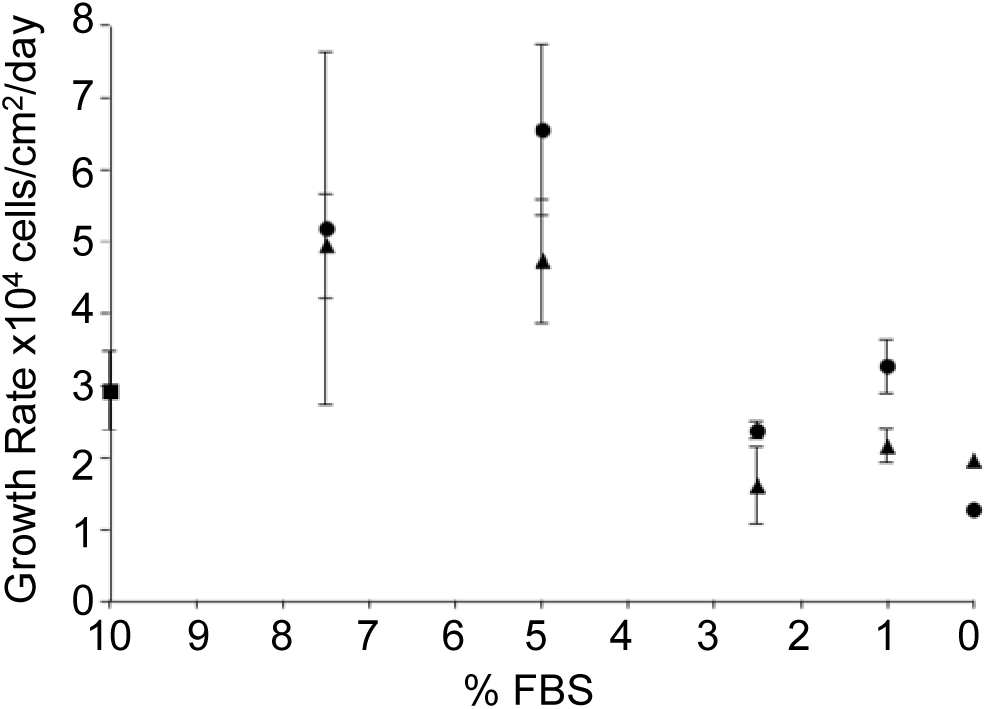
Average growth rates of CV-1 cells during stepwise adaption to serum-free media. Starting at 10% FBS DMEM (square) cells were harvested and re-plated in mixtures of serum-containing and serum-free medium to give the serially decreasing overall serum concentration indicated by the X-axis. VP-SFM (triangles) or OptiPRO SFM (circles) brands of serum free medium were used. Growth rates were determined as detailed in Methods.

### Cell Growth Kinetics in T flasks

Average cell growth rate (cells/cm^2^/day) was calculated using Equation 1,

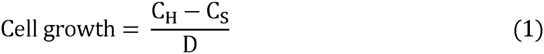

-where C_H_ is total cell density (cells/cm^2^) at harvest; C_S_ is the total cell density (cells/cm^2^) at seeding and D is culture duration (days). Cell Recovery Ratio (CRR) under complete serum free conditions was calculated using Equation 2.

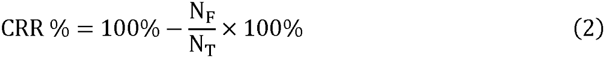

-where N_F_ is total number of detached cells 24 hours post seeding; N_T_ is total number of cells seeded (cells/cm^2^). Specific growth rate, μ(h^−1^) was based on Equation 3.

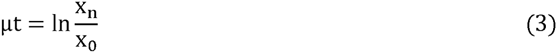

Where x_0_ is starting total cell concentration per mL; t is the time of sampling in hours; x is total cell concentration per mL after t hours. According to Eq. (1), a plot of ln x versus time gives a straight-line plot with μ as the slope. Cell doubling time, DT (hours) was calculated using Equation 4, where μ_max_ is the maximum specific growth rate during the exponential phase, hour^−1^.

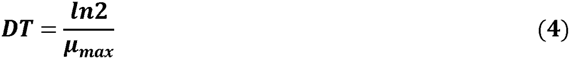

### Virus Propagation

A single virus stock was used throughout this work. A titre of 6.68×10^8^ PFU/mL was determined for this stock using the procedures described below and with CV-1 cells used for titration. Aliquots for infection were diluted with the required volume of growth media. Virus solutions were added drop wise to cells at a density of 3-5×10^5^ cells/well in a 6-well plate at a multiplicity of infection (MOI) of 0.1. After the indicated time period, infected ‘propagator’ cells were harvested using a cell scraper for virus release by cell disruption.

### Virus release from propagating cells

Suspensions of cells infected for virus propagation were frozen in a -80°C freezer for 30 minutes, thawed in a 37°C water bath for 4 minutes and vortexed for 10 seconds. This freeze-thaw-vortex cycle was repeated three times and the resultant disruptate containing cell debris and released virus particles used for virus titration with no further purification.

### Virus infection of target cells for titration

Median tissue culture infective dose (TCID_50_) was determined using CV-1 cells as target cells. Disruptates, containing cell debris and viruses, were serially diluted in 96-well plates containing cells adapted to, and grown in, 5% FBS DMEM unless otherwise stated. Cytopathic effect (CPE) was scored by light microscopy six days post infection. The Reed-Muench procedure (1938) was used to calculate TCID_50_ values, which were converted to PFU/cell using Equation 5.

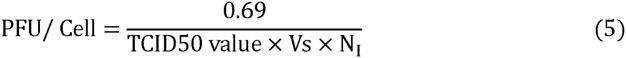

-where Vs is the volume of sample used to infect the first row of the 96-well titration plates, mL; N_I_ is number of cells at infection.

### Growth on microcarriers of CV-1 cells adapted to OptiPRO medium

#### Pre-treatment of vessels and microcarriers

Cultivation of OptiPRO-adapted CV-1 cells adhered to Cytodex-1 microcarriers (GE Healthcare, Westborough, USA) was performed using a Techne MCS-104L 250mL spinner flask setup (Bibby Scientific Ltd, Staffordshire, UK). All procedures were performed in a Level 1 laminar flow biological safety cabinet (BSC) unless otherwise stated. Spinner flasks were first prepared for use by siliconisation of the flask interior surface and impellers using Sigmacote (SL2, Sigma-Aldrich, USA) in accordance with manufacturer’s instructions. When the siliconisation procedure was complete the impeller system was assembled within the 250mL spinner flask and autoclaved using a cycle of 20 minutes at 121°C.

A 0.5L pyrex Duran bottle was also coated with silicon using the Sigmacote and following manufacturer’s instructions. Briefly, 100mL of Sigmacote were poured into to the 0.5L pyrex Duran bottle which was swirled until all the interior surface received a covering of Sigmacote. Remaining Sigmacote was decanted and the bottle was autoclaved, dried in a fume hood overnight then rinsed with Milli-Q water to remove any siliconisation by-products before use.

The required mass of Cytodex-1 microcarriers (17-0448-01, GE Healthcare, Sweden) was added to the siliconised Pyrex Duran bottle. For every gram of dry microcarriers 100mL of pH 7.4 PBS, free of calcium and magnesium ions (Life Technologies, Paisley, UK), was added. The bottle was left to stand over night to achieve complete swelling and hydration of the microcarriers. The next day the PBS used to hydrate the microcarriers was gently decanted and replaced with 50mL fresh PBS for every gram of wet microcarriers. The microcarrier slurry was then sterilized by autoclaving (121 °C, 20 min). Prior to use, the sterilized microcarriers were allowed to cool and settle. Upon cooling the supernatant was gently decanted and replaced with 50 mL fresh OptiPRO medium for every gram of wet microcarriers. This OptiPRO/microcarrier slurry was allowed to settle and the OptiPRO supernatant was gently decanted then replaced with 100 mL fresh OptiPRO for every gram of wet microcarriers.

#### Mixing cells and microcarriers

30mL of the microcarrier / media slurry was transferred to the spinner flask. 70mL of a suspension of OptiPRO-adapted CV-1 cells in OptiPRO was then added to the spinner flask to achieve a microcarrier concentration of 3g/L and a cell concentration of 3.4x10^5^ cells/mL, corresponding to 26 cells / microcarrier. The spinner flask was then placed on the MCS-104L stirrer device inside an MCO-19AIC incubator (Sanyo, Gunma, Japan) at 37°C and 5% CO_2_. The mixture of cells and microcarriers was subjected to intermittent agitation at 30 RPM for 30 seconds every 30 minutes for the first two days of cultivation after which continuous 30 RPM agitation was used for the third day and then 35 RPM for the remainder of the cultivation experiment. Approximately 70% of the volume of culture medium was replaced with fresh OptiPRO every 24 hours by terminating agitation; allowing microcarriers to settle decanting supernatant and adding fresh OptiPRO.

#### Removing CV-1 cell samples during microcarrier-based cultivation

The MCS-104L stirrer device was switched off and transferred, with the spinner flask, from the incubator to the BSC. The stirrer was set to agitate the spinner flask at 60 RPM and the cap from one side-arm port of the spinner flask was removed to allow withdrawal of a 0.2mL sample from the culture. The MCS-104L stirrer device was then switched off and the entire setup returned to the incubator where incubation and agitation were resumed.

#### Counting CV-1 cells during cultivation using microcarriers

Samples taken as above were typically transferred to a 1.5mL Eppendorf tube and washed with 1mL PBS. Microcarriers were allowed to settle and 1ml supernatant was gently decanted; then a further 1mL PBS wash performed. After the final decanting of supernatant 0.2 mL aqueous crystal violet solution (0.1% w/v Crystal Violet, 0.1M citric acid, 0.1% v/v Triton X-100) was added to the microcarriers slurry and the mixed by pipetting up and down 25 times before the Eppendorf tube was transferred to an incubator set at 37°C / 5% CO_2_ for 1.5 hours. This treatment causes cells to lyse and release stained nuclei. Typically the solution was diluted by addition of PBS. Released nuclei were counted using an Improved Neubauer haemocytometer (1080346, Heinz Herenz Medizinalbedarf GmbH, Hamburg, Germany) as an indicator of cell numbers.

## Results and Discussion

### Considerations for VACV genome refactoring and BioBrick™-based VACV plasmid tools

Industrial synthetic biology seeks to go beyond conventional bioprocessing and design host organisms (‘chassis’) with predictable process behaviour. In the case of VACV biomanufacturing, a first step toward such a synthetic biology platform would involve refactoring the VACV genome for improved safety and manipulability. Chan et al. (2005) proposed a refactored version of the T7 bacteriophage genome with a major design goal being to enable the precise and independent manipulation of discrete elements, such as open reading frames (ORFs). This goal was met by repositioning unique restriction sites naturally present in the T7 genome and introducing new unique sites, previously absent from the genome. We analysed the VACV Lister strain genome (Figure 1A) and located 6 unique restriction sites, cut by enzymes SanDI, SbfI, NotI, SgrAI, SmaI and ApaI in that order. AscI, DpnI, FseI, RsrII, SfiI, SgfI and SrfI sites are absent from the VACV Lister strain genome so may be candidates for addition by refactoring.

**Figure 1.**
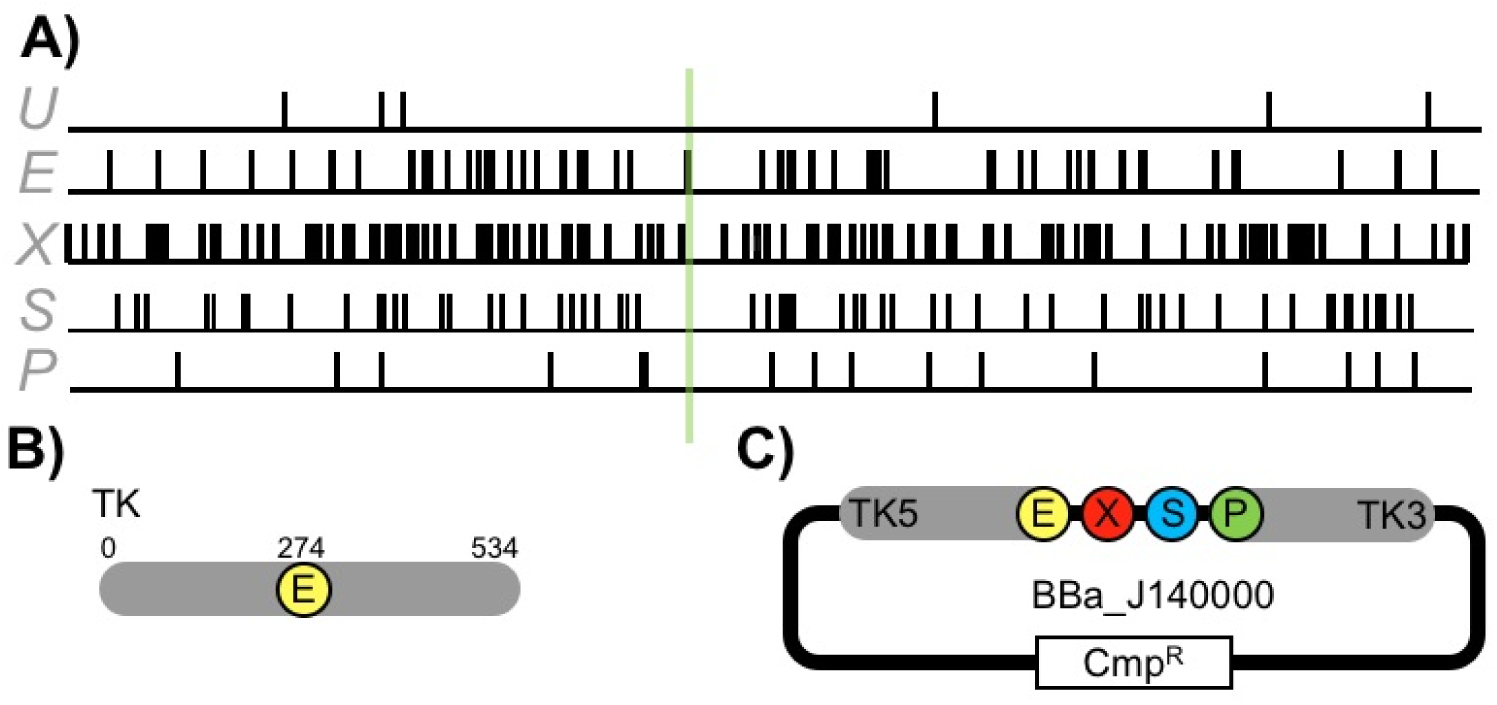
Considerations for refactoring the Vaccinia virus Lister strain genome and design of a BioBrick™-formatted Vaccinia DNA tool. Panel A: restriction sites mapped as vertical lines joining a horizontal line representing, from left to right in 5’ to 3’ direction, the 189.4kb Lister strain VACV genome (GenBank: DQ121394.1). Maps, drawn using ApE software (Davis MA, U. Utah, USA), indicate, from the top, the locations of 6 unique sites (U), 53 EcoRI (E), 125 XbaI (X), 61 SpeI (S) and 17 PstI (P) sites in the Lister strain VACV genome. Green line indicates locus of Thymidine Kinase (TK) gene (GenBank: ABD52560.1). Panel B: location of lone Eco R1 site within 534bp TK ORF. Panel C: Design of a BioBrick™-formatted plasmid tool (BBa_J140000) for VACV genome editing, incorporating a selection marker (open box), and the E, X, S, P BioBrick™ cloning site flanked by 5’ and 3’ ends of the TK locus for homologous recombination.

We also indicated the location of the EcoRI, XbaI, SpeI and PstI sites in the Lister strain VACV genome (Figure 1A) as these sites define the BioBrick™ DNA fragment assembly standard (Shetty et al. 2008) widely used in the synthetic biology community and beyond. The six-fold variation in prevalence of these sites in the VACV genome serves to indicate the scale of refactoring that can be required to render natural DNA more compatible with standardised synthetic biology formats even for relatively small viral genomes.

The VACV genome is conventionally edited by parallel viral infection and plasmid transfection of mammalian cells. The plasmids contain regions of homology with a VACV genomic locus and this directs targeted recombination that results in progeny virions with edited genomes. Typically the locus encoding the ORF for thymidine kinase (TK) is used to target insertion of transgenes (Byrd and Hruby, 2004) as its disruption does not compromise virus replication in cells commonly used for tissue culture (Figure 1B). The TK ORF contains a lone EcoRI restriction site at its approximate mid-point (Figure 1B). The location of this EcoRI site suggests a convenient point for splitting the TK and positioning it either side of the EcoRI, XbaI, SpeI and PstI BioBrick™ cloning site in a BioBrick™-compatible VACV genome editing tool (Figure 1C). BioBrick™-based tools for virus design have been developed previously, such as BBa_K404129 that encodes a transgene expression cassette designed to be encapsidated by adeno-associated virus. Our proposed design, BBa_J140000, opens up the VACV genome to exploration using all components of the Registry of Standard Biological Parts (International Genetically Engineered Machines, Boston, USA).

### Adaptation of CV-1 cells to serum free growth media

CV-1 is a continuous cell line derived from *Cercopithecus aethiops* African green monkey kidneys by Jensen (1964). It is susceptible to several viruses including VACV and has been widely used for virus titration (Cho et al., 1970; Hiley et al., 2010) but is previously unused for VACV propagation. Given its proven track record of both safe cultivation and efficient viral infection we chose CV-1 for this preliminary investigation. We sourced CV-1 cells from ATCC (product CCL-70™) and considered them as already adapted to growth in 5% FBS DMEM.

Cells were grown in a rich medium, 10% FBS DMEM, before stepwise adaptation to growth in the VP-SFM and OptiPRO brands of serum-free medium. The serum content (v/v) of the growth medium mix used during each round of adaptation was lowered to 7.5%, 2.5%, 1% and finally 0%. Figure 2 shows the average growth rates observed when cells were first challenged with the decreased serum content media mix. Average CV-1 cell growth rate in 10% FBS DMEM was 2.93±0.54 x10^4^ cells/cm^2^/day. Average growth rates in 7.5% and 5% FBS media mixes, for both VP-SFM and OptiPRO, were increased compared to 10% FBS DMEM. Only 0% serum, pure VP-SFM or OptiPRO, resulted in initial growth rates lower than that for 10% FBS DMEM, with 1.97x10^4^ cells/cm^2^/day and 1.29x10^4^ cells/cm^2^/day respectively. Cells were then grown in VP-SFM or OptiPRO for another four passages before being considered as fully adapted to serum-free media.

### Growth profiles of CV-1 cells adapted to serum-free media

Cells adapted to 5% FBS DMEM, VP-SFM and OptiPRO were seeded at 5x10^4^ cells/25cm^2^ in T-25 flask and cell densities measured every 24 hours for 350 hours to observe lag, exponential and stationary phases (Figure 3) and determine growth rate (Table 1). For cells adapted to 5% FBS DMEM (Figure 3A), total cell numbers decreased over the first 24 hours post-seeding. Growth increased after this with a specific growth rate of 0.035 h^−1^ observed, corresponding to a doubling time of 20.1 hours. This is comparable to the doubling time of 22 hours reported by Hagedorn et al. (1985) for CV-1 cells grown in medium with 5% v/v foetal calf serum (FCS). Cell growth slowed after 168 hours, at a saturation density of 4.52x10^6^ cells/25cm^2^. Saturation density of adherent cells on a solid surface is a potential indicator of microcarrier growth performance.

**Figure 3.**
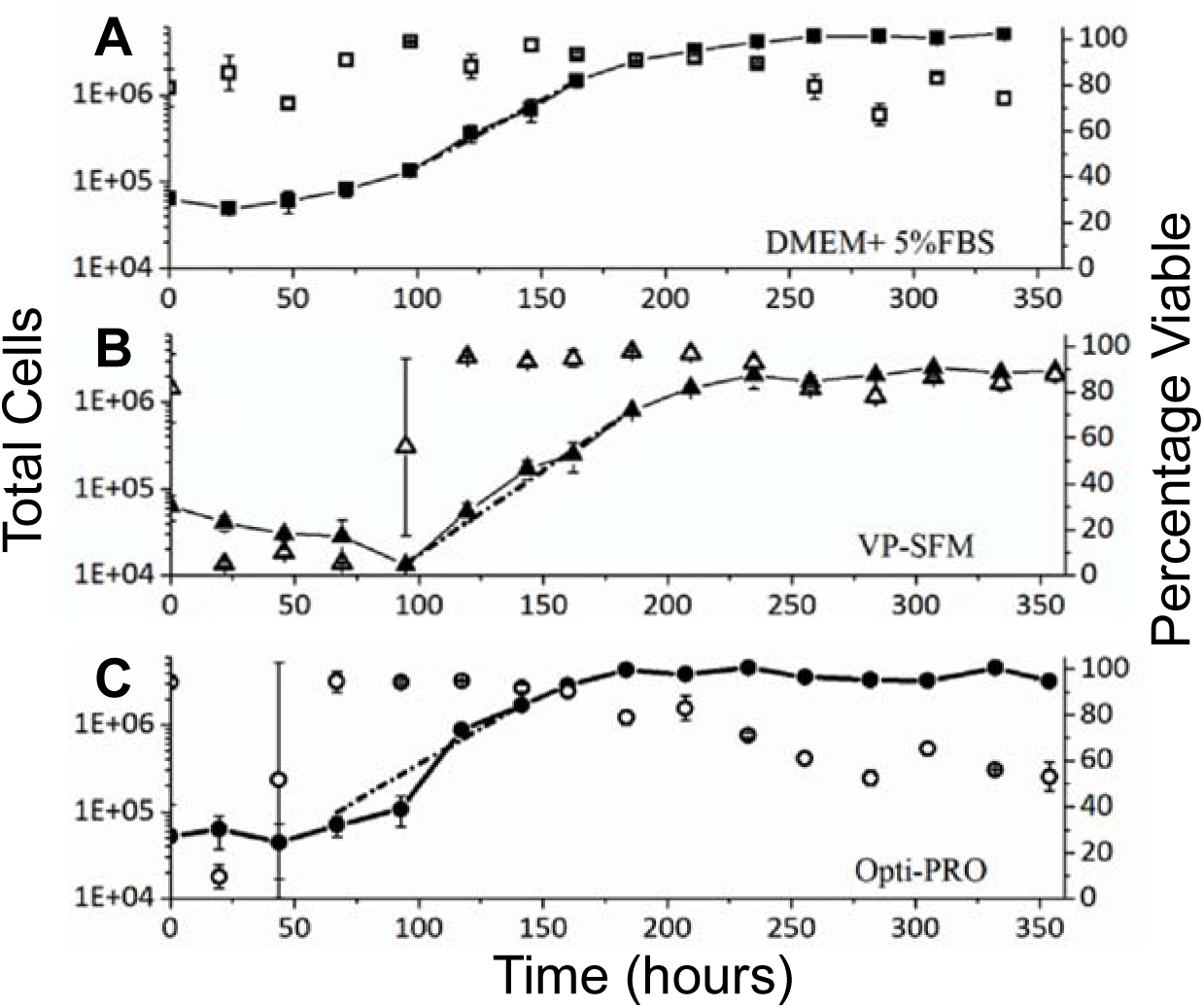
Growth of CV-1 cells adapted to serum-free media. Cells adapted to, and grown in, A) 5% FBS DMEM (squares), B) VP-SFM (triangles) and C) OptiPRO (circles) were seeded into T-25 flasks at the density indicated and their growth (closed symbols) and viability (open symbols) followed over 350 hours. Total cells were detached by trypsinisation, counted and reseeded in fresh media at each time point, with 1-5% of material discarded after cell counting and viability assessment. Dashed line indicates the linear regression of cell growth during exponential phase used to calculate growth rates provided in Table 1. Error bars indicate standard deviation over n=2 biological repeats.

**Table 1.**
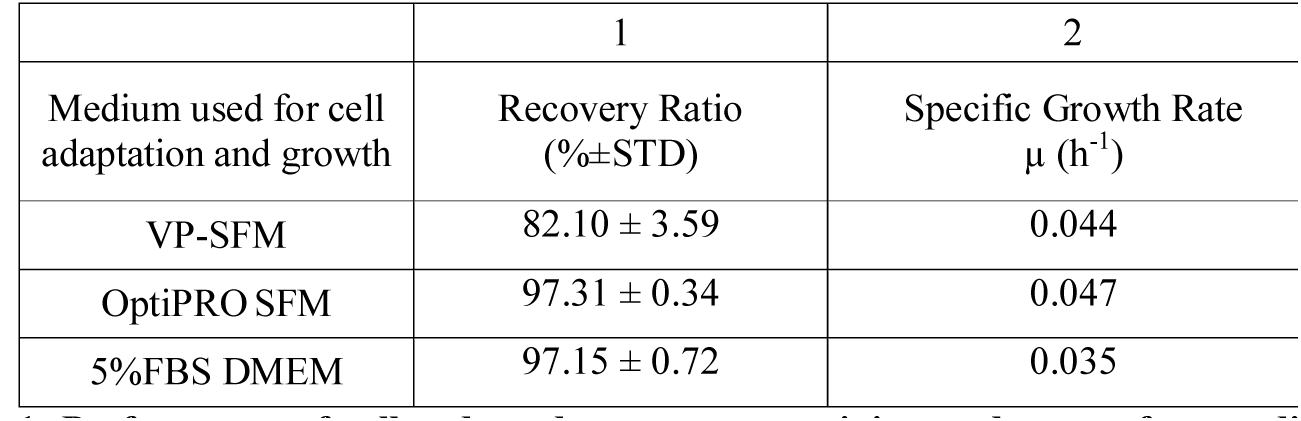
Performance of cells adapted to serum-containing and serum-free media types. Summary of performance data for cells adapted to growth in the different media types. Column 1 shows recovery from cryopreservation (recovery ratio). Column 2 provided growth rates of cells after adaptation to the indicated media type.

| Medium used for cell adaptation and growth | 1 | 2 |
| --- | --- | --- |
| | Recovery Ratio (%±STD) | Specific Growth Rate $\mu$ (h<sup>-1</sup>) |
| VP-SFM | 82.10 ± 3.59 | 0.044 |
| OptiPRO SFM | 97.31 ± 0.34 | 0.047 |
| 5%FBS DMEM | 97.15 ± 0.72 | 0.035 |

Cells adapted to VP-SFM (Figure 3B), decreased rapidly in number over the first 95 hours post-seeding, with viability as low as 4.9%. Cells then entered exponential growth, achieving 90% viability and a specific growth rate of 0.044 h^−1^ (15.9 hours doubling time). Cells entered stationary phase 235 hours post-seeding with saturation density of 2.15x10^6^ cells/25cm^2^.

Growth of cells adapted to OptiPRO (Figure 3C) lagged over the first 67 hours post-seeding then grew exponentially, with a specific growth rate of 0.047 h^−1^ (14.8 hours doubling time). These cells reached stationary phase at 192 hours post-seeding, at a saturation density of 3.84x10^6^ cells/25cm^2^. Notably, a significant decrease in viability was observed 24 hour post-seeding. This increased to >90% at 67 hours, remained above 90% until 200 (160) hours then declined to <60% at 354 hours post-seeding.

Compared to cells adapted to grow in 5% FBS DMEM, cells adapted to VP-SFM and OptiPRO had lower saturation densities and significant drops either in cell number or viability in the first 96 hours post-seeding. The absence of specific growth factors (Todarro et al. 1965, Dulbecco et al. 1973, Vogel et al. 1980) or nutrients from certain serum-free media formulations may result in reduced shear resistance in mammalian cells (El-Ensahsy et al., 2009). This could explain the extended lag phase growth of cells adapted to VP-SFM and OptiPRO compared to those adapted to 5% FBS DMEM (Figure 3). VP-SFM and OptiPRO-adapted cells may have required a longer time period to recover from shear experienced over multiple rounds of detachment, resuspension and re-seeding during adaptation (Figure 2).

### CV-1 cell robustness to cryopreservation

Stable and reliable recovery from cryopreservation is a critical attribute of mammalian cells used for industrial production of biotherapeutics. Recovery ratio provides an indication of the effectiveness of a given formulation of cryopreservant media for storing cells under liquid nitrogen. Cells adapted to growth in 5% FBS DMEM, VP-SFM and OptiPRO were resuspended in cryopreservant media, as detailed in the Methods section above, containing methylcellulose as a protective agent only for CV-1 cells previously adapted to VP-SFM and OptiPRO (Waymouth et al. 1976). After storage in liquid nitrogen for six months cells were revived and recovery ratios determined (Table 1). Recovery ratios of ≈97% were measured for cells grown in, and adapted to, both OptiPRO and 5% FBS DMEM. Cells adapted to VP-SFM had the lowest recovery ratio of ≈82%.

### Vaccinia virus production by CV-1 cells adapted to grow in serum-free media

For production of VACV strain TSI-GSD-241 using MRC-5 cells for propagation, Wu et al. (2005) reported virus productivity of 77 PFU/cell when the cells were grown in 20% FBS DMEM and infected at an MOI of 0.1. We used a recombinant Lister strain VACV, VACVL-15 RFP, encoding a red fluorescent protein expression cassette payload.

At an MOI of 0.1 we infected CV-1 cells (as ‘propagators’) adapted to, and cultivated in, 5% FBS DMEM, VP-SFM or OptiPRO. Infected cells were incubated at 37°C with 5% CO_2_ for 24, 48 and 72 hours post-infection after which virus was released and virus titre measured (Figure 4) by infection of CV-1 cells adapted to, and cultivated in, 5% FBS DMEM as ‘targets’ cells. At 72 hour post infection ‘propagator’ cells adapted to growth in OptiPRO achieved a titre of 352 PFU/cell, 4.6 times higher than the titre achieved by cells adapted to 5% FBS DMEM and 2.6 times higher than the titre achieved by cells adapted to VP-SFM.

**Figure 4.**
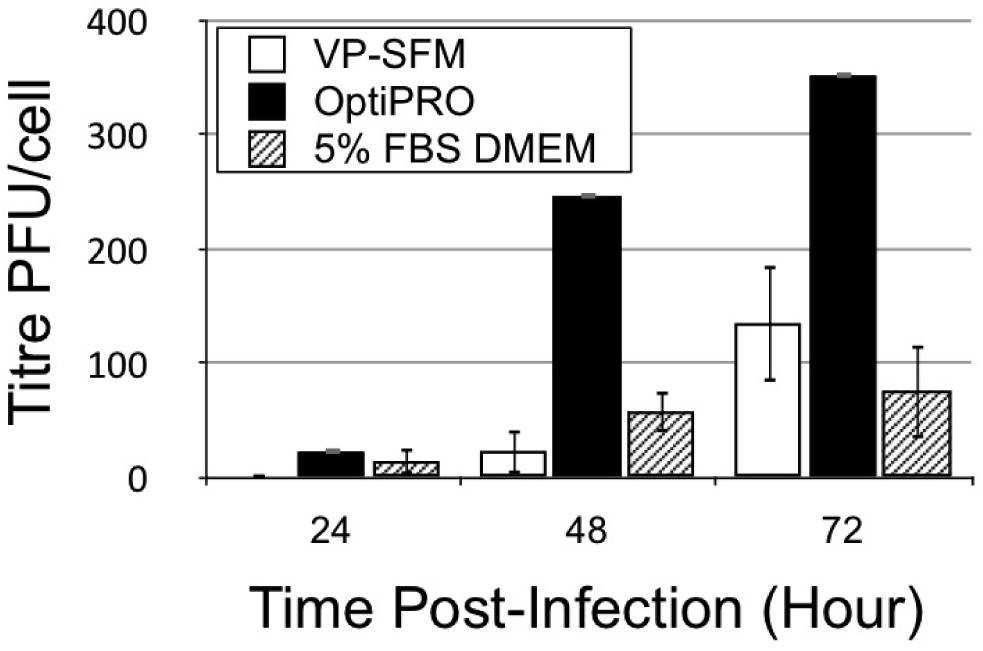
Viral productivity of CV-1 cells adapted for growth in serum free media. Vira l titre productivity of CV-1 cells adapted for growth in VP-SFM (open bar), OptiPRO (black bars) and 5% FBS DMEM (striped bars). Cells at 80-95% confluence were infected with VACVL-15 RFP at MOI=0.1 and virus liberated for titration at indicated times post-infection. Error bars indicate standard deviation over n=3 biological repeats.

### Influence of media type and cell provenance on viral titre performance

It is not evident from Figure 4 whether the enhanced titre performance of OptiPRO-adapted cells results from OptiPRO favouring virus infection events or OptiPRO exerting a selective pressure that favours CV-1 cells capable of high virus productivity. Furthermore, compatibility with a notional synthetic biology production platform for VACV manufacture would require multiple iterations of entirely serum-free propagation.

As such we repeated the OptiPRO, 72 hour post-infection harvest time experiment of Figure 4 alongside two comparator experiments in an attempt to determine both the likely causative factors for the increased titre observed in Figure 5 and the relative efficiency of an entirely serum-free round of propagation. Table 2 summarises our approach; cells adapted for growth in OptiPRO were grown to 95% confluence then washed twice with PBS before immersion either again in OptiPRO (Figure 5A and 5C) or 5% FBS DMEM (Figure 5B), immediately prior to infection. For titration, OptiPRO-adapted cells in the presence of OptiPRO (Figure 5A) and 5% FBS DMEM-adapted cells in the presence of 5% FBS DMEM (Figure 5B and 5C) were used as ‘targets’ for titre measurement.

**Figure 5.**
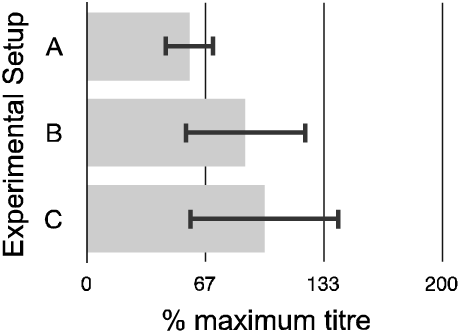
Influence of media and cell type on viral titre performance. As set out in Table 2, CV-1 cells were washed in PBS and re-immersed either in OptiPRO or 5% FBS DMEM, infected with VACV at MOI=0.1 and 72 hours post-infection progeny virus used to infect cells adapted to growth in OptiPRO in the presence of OptiPRO (Experimental setup A) or 5% FBS DMEM (Experimental setup B), or cells adapted to growth in 5% FBS DMEM in the presence of 5% FBS DMEM (Experimental setup C). Error bars indicate standard deviation over n=3 biological repeats. As Experimental setup C was a repeat of the experiment performed to generate the data in Figure 4 (72 virus harvest 72 hours post-infection), the tire achieved with Experimental setup C was set as the 100% level.

**Table 2.** Dissecting effect of growth medium and cell-adaptation on viral infection and production. <u>Experiment setup A</u>: CV-1 cells adapted to growth in OptiPRO were washed in PBS, re-immersed in OptiPRO and infected with VACV at MOI=0.1. 72 hours post-infection, progeny virus from these cells was used to infect CV-1 cells adapted to growth in OptiPRO, in the presence of OptiPRO. <u>Experiment setup B</u>: CV-1 cells adapted to growth in OptiPRO were washed in PBS, immersed in 5% FBS DMEM and infected with VACV at MOI=0.1. 72 hours post-infection, progeny virus from these cells was used to infect CV-1 cells adapted to growth in 5% FBS DMEM, in the presence of 5% FBS DMEM. <u>Experiment setup C</u>: CV-1 cells adapted to growth in OptiPRO were washed in PBS, re-immersed in OptiPRO and infected with VACV at MOI=0.1. 72 hours post-infection, progeny virus from these cells was used to infect CV-1 cells adapted to growth in 5% FBS DMEM, in the presence of 5% FBS DMEM. Data generated from these experiments was plotted in Figure 5. *Propagator: cells that are infected with virus for the purpose of harvesting virus particles from those cells. **Target: cells that have been infected with virus in order to establish a TCID_50_ as an indication of the titre of the virus solution.

| Experiment | A | B | C |
| --- | --- | --- | --- |
| **Propagator cell\*** | Cells adapted to growth in OptiPRO | Cells adapted to growth in OptiPRO | Cells adapted to growth in OptiPRO |
| **Media present during Propagator cell infection** | OptiPRO | 5% FBS DMEM | OptiPRO |
| **Target cell\*\*** | Cells adapted to growth in OptiPRO | Cells adapted to growth in 5% FBS DMEM | Cells adapted to growth in 5% FBS DMEM |
| **Media present during Target cell infection** | OptiPRO | 5% FBS DMEM | 5% FBS DMEM |

Experiment C (Table 2) is a straight repeat of the conditions used in Figure 4 (data in black bars, harvest 72 hours post-infection) so the resultant titre was set as the 100% level for comparison with Experiments A and B (see Table 2, Figure 5A and Figure 5B). If the presence of OptiPRO media enhances VACV infection of CV-1 cells, then Experiment A could be expected to increase the titre achieved by Experiment C (see Table 2, Figure 5A and Figure 5C). This is not the case, with Experiment A yielding at best the same titre performance as Experiment C. If the presence of 5% DMEM enhances VACV infection of CV-1 cells, then Experiment B could be expected to increase the titre achieved by Experiment C (see Table 2, Figure 5B and Figure 5C). This is not the case, with Experiment B yielding at best the same titre performance as Experiment C.

Taken together, observations from Figures 4 and 5 are consistent with the enhanced titre observed for OptiPRO-adapted cells being due to the process of adaptation to OptiPRO media also exerting a post-infection phenotype of increased virus productivity. They also indicate that entirely serum-free rounds of VACV propagation, such as those likely to define industrial synthetic biology platforms, yield comparable titre performance to serum-containing processes and so are feasible.

### CV-1 cell cultivation using OptiPRO and microcarriers

We sought to determine if CV-1 cells adapted to OptiPRO could be cultivated using microcarriers (Figure 6). We attempted to cultivate cells using Cytodex-1 microcarriers and a Techne MCS-104L 250mL spinner flask setup (Bibby Scientific Ltd, Staffordshire, UK) in which microcarrier suspensions are agitated by bulb–shaped glass impellers driven directly by a magnetic base. OptiPRO-adapted CV-1 cells were also able to grow on micocarriers when approximately 70% of OptiPRO mediumwas changed daily and showed reduced growth when the OptiPRO made was unchanged over 270 hours of cultivation (Figure 6).

**Figure 6.**
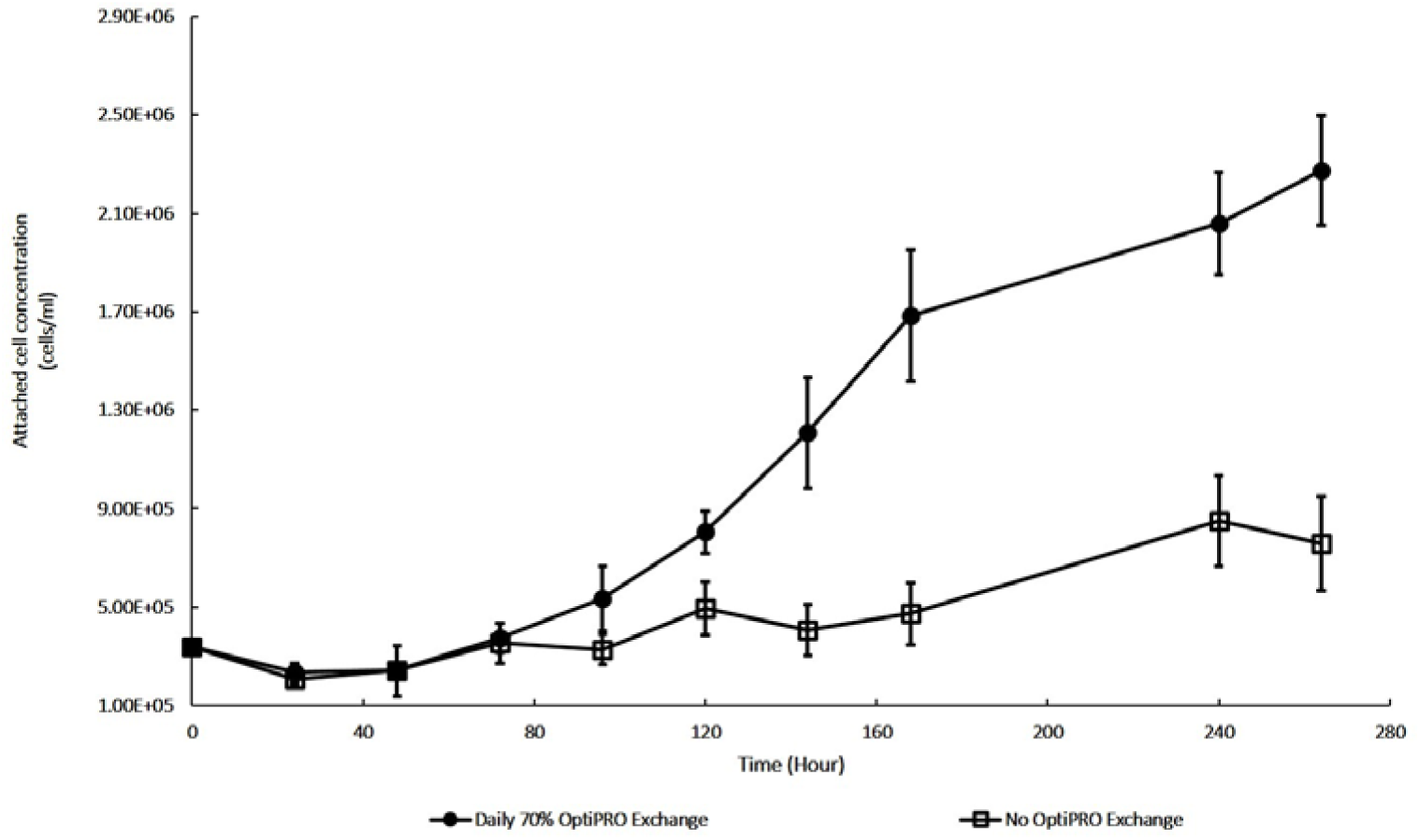
Cultivation of CV-1 cells adapted to OptiPRO media on microcarriers. Cells adapted for growth in OptiPRO were mixed with Cytodex-1 microcarriers at a starting ration of 26:1 in 100mL OptiPRO and agitated at 30-35 RPM over the indicated period. Cell concentrations were plotted for cultivation in which either 70% of the OptiPRO was changed every 24 hours (black circles) or OptiPRO was unchanged throughout (open squares). Error bars indicate standard deviation over n=3 biological repeats.

In the case of T-flask cultivation, VACV propagation involves two significant factors: virus number and cell numbers, which are summarised by the MOI. By contrast microcarrier-based VACV propagation presents three major factors; viruses, cells and microcarriers, and as such represents a complex investigation to identify productivity optima, such as those reported by Monath et al. (2004) for production of VACV from Vero cells grown using microcarriers and Bleckwernn et al at (2005) for VACV production from HeLa cells. Such an investigation falls outside the scope of this study, which is to indicate the broad feasibility of the steps likely to define a future synthetic biology platform for VACV production.

## Conclusions

We appraised the VACV genome in light of refactoring considerations and proposed a BioBrick™ plasmid for standardised VACV genome insertion. CV-1 cells adapted for growth in OptiPRO serum free medium exhibited elevated titre performance when grown using static culture and showed promising growth characteristics on Cytodex-1 microcarriers. Mechanisms underlying the elevated titre are unclear but may result from selective pressure exerted by the adaptation process acting also to select for an unintended phenotype.

Overall these results are consistent with the assertion that a standardised, serum-free, microcarrier-based synthetic biology platform for production of VACV is feasible. Cultivation in suspensions is inherently more scalable than cultivation on planar surfaces so further scale up of the platform proposed here should be investigated. Future work should include an investigation of the optimal conditions for VACV production from CV-1 cells grown on microcarriers, including the ratio of infecting virus to cells and microcarriers. Characterisation of virus quality should also be performed to assess factors such as the percentage of total virus particles that are plaque-forming as opposed to inactive.

## References

The European Agency for the Evaluation of Medical Products (2002) Note for guidance on the development of Vaccinia virus based vaccines against smallpox. European Medicines Agency. http://www.ema.europa.eu/docs/en_GB/document_library/Scientific_guideline/2009/09/WC500003900.pdf Accessed 18 October 2015

Barrett PN, Mundt W, Kistner O, Howard MK. (2009) Vero cell platform in vaccine production: moving towards cell culture-based viral vaccines. Expert Rev Vaccines 8(5):607–18.

Bleckwenn NA, Bentley WE, Shiloach J. (2005) Evaluation of production parameters with the vaccinia virus expression system using microcarrier attached HeLa cells. Biotechnol Prog 21(2):554–61.

Byrd CM, Hruby DE. (2004) Construction of recombinant vaccinia virus: cloning into the thymidine kinase locus. Methods Mol Biol 269:31–40.

Chan LY, Kosuri S, Endy D. (2005) Refactoring bacteriophage T7. Mol Syst Biol 2005.0018.

Cho CT, Locke T, Wenner HA. (1970) Viremia and Virus Measurements of RabbitPox in CV-1 Cells. Appl Microbiol 19(5):791–794

Dymond JS, Richardson SM, Coombes CE et al (2011) Synthetic chromosome arms function in yeast and generate phenotypic diversity by design. Nature 477(7365):471–476.

EL-Ensahsy HA, Abdeen A, Abdeen S. (2009) Serum concentration effects on the kinetics and metabolism of HeLa-S3 cell growth and cell adaptability for successful proliferation in serum free medium. World Appl Sci J 6 (5): 608–615.

Fenner F, Isao A, Henderson DA, Jezek Z, Ladnyi ID. (1988) The history of smallpox and its spread around the world. In: Fenner F, Isao A, Henderson DA, Isao A, Jezek Z, Ladnyi ID (eds) Smallpox and its eradication, World Health Organization, Geneva, pp 209–244.

Garcel A, Crance JM, Drillien R, Garin D, Favier AL. (2007) Genomic sequence of a clonal isolate of the vaccinia virus Lister strain employed for smallpox vaccination in France and its comparison to other orthopoxviruses. J Gen Vir 88:1906–1916.

Guse K, Cerullo V, Hemminki A. (2011) Oncolytic vaccinia virus for the treatment of cancer. Expert Opin Biol Ther 11(5):595–608.

Hagedorn R, Thielmann HW, Fischer H. (1985) SOS-Type functions in mammalian cells enhanced reactivation of UV-irradiated SV 40 in UV-irradiated CV-1 Cells. J Cancer Res Clin Oncol 109(2):89–92.

Hiley CT, Yuan M, Lemoine NR, Wang Y. (2010) Lister strain vaccinia virus, a potential therapeutic vector targeting hypoxic tumours. Gene Ther 17(2):281–7.

Hruby DE. (1990) Vaccinia virus vectors: new strategies for producing recombinant vaccines. Clin Microbiol Rev 3(2):153–170.

Jensen FC. (1964) Infection of human and simian tissue cultures with Roussarcoma virus. Proc Nat Acad Sci U S A 52:53–59.

Kononenko AV, Lee NCO, Liskovykh M et al (2015) Generation of a conditionally self-eliminating HAC gene delivery vector through incorporation of a tTA^VP64^ expression cassette. Nucl Acids Res. doi: 10.1093/nar/gkv124

Lee SS, Eisenlohr LC, McCue PA, Mastrangelo MJ, Lattime EC. (1994) Intravesical gene therapy: in vivo gene transfer using recombinant vaccinia virus vectors. Cancer Res 54(13):3325–8.

Mackett M, Smith GL (1986) Vaccinia virus expression vectors. J Gen Virol 67 (Pt 10):2067–82.

Mayrhofer J, Coulibaly S, Hessel A et al (2009) Nonreplicating Vaccinia virus vectors expressing the H5 Influenza virus hemagglutinin produced in modified Vero cells induce robust protection. J Virol 83(10):5192–5203.

Monath TP, Caldwell JR, Mundt W et al (2004) ACAM2000 clonal Vero cell culture vaccinia virus (New York City Board of Health strain)–a second-generation smallpox vaccine for biological defence. Int J Infect Dis 8 Suppl 2:S31–44.

Reed LJ, Muench H. (1938) A simple method of estimating fifty percent endpoints. The American Journal of Hygiene 27(3): 493–497.

Schweneker M, Lukassen S, Späth M, Wolferstätter M, Babel E, Brinkmann K, Wielert U, Chaplin P, Suter M, Hausmann J. (2012) The Vaccinia Virus O1 Protein Is Required for Sustained Activation of Extracellular Signal-Regulated Kinase 1/2 and Promotes Viral Virulence. J Virol. 2012 Feb; 86(4): 2323–2336.

Shetty RP, Endy D, Knight TF Jr. (2008) Engineering BioBrick™ vectors from BioBrick™ parts. J Biol Eng 2:5.

Steimer KS, Packard R, Holden D, Klagsbrun M. (1981). The serum-free growth of cultured cells in bovine colostrum and in milk obtained later in the lactation period. J Cell Physiol. 109(2):223–34.

Timiryasova TM, Li J, Chen B et al (1999) Antitumor effect of vaccinia virus in glioma model. Oncol Res 11(3):133–144.

Todaro GJ, Lazar GK, Green H. (1965) The limitation of cell division in a contact-inhibited mammalian cell line. J Cell Physiol 66:325–333.

Tysome JR, Wang P, Alusi G et al (2011) Lister vaccine strain of vaccinia virus armed with the endostatin-angiostatin fusion gene: an oncolytic virus superior to dl1520 (ONYX-015) for human head and neck cancer. Hum Gene Ther 22(9):1101–8.

Vogel A, Ross R, Raines E. (1980) Role of serum components in density-dependent inhibition of growth in 3T3 cells: platelet-derived growth factor is the major determinant of saturation density. J. Cell Biol 85(2):377–85.

Waymouth C, Vamum DS. (1976) Simple freezing procedure for storage in serum-free media of cultured and tumor cells of mouse. TCA manual 2(1):311–313.

Wu F, Reddy K, Nadeau I, Gilly J, Terpening S, Clanton DJ. (2005) Optimization of a MRC-5 cell culture process for the production of a smallpox vaccine. Cytotechnology 49 95–107.

Ye H, Fussenegger M. (2014) Synthetic therapeutic gene circuits in mammalian cells. FEBS Lett 588(15):2537–44.

Zhang Q, Yu YA, Wang E et al (2007) Eradication of solid human breast tumors in nude ice with an intravenously injected light-emitting oncolytic vaccinia virus. Cancer Res 67(20):10038–46.

